# Delay-related activity in marmoset prefrontal cortex

**DOI:** 10.1101/2022.04.04.487026

**Authors:** Raymond Ka Wong, Janahan Selvanayagam, Kevin Johnston, Stefan Everling

**Author notes:** Corresponding author: Stefan Everling, Robarts Research Institute, University of Western Ontario, London, Ontario, Canada. **Email:**. **Author Contributions** RW performed experiments, analysed data, prepared figures and wrote the manuscript. JS assisted in performing experiments and data analysis. KJ assisted in surgeries and data analysis. SE designed experiments and performed surgeries. All authors edited the manuscript and SE approved the final version.

## Abstract

Persistent delay-period activity in prefrontal cortex (PFC) has long been regarded as a neural signature of working memory (WM). Electrophysiological investigations in macaque PFC have provided much insight into WM mechanisms, however a barrier to understanding is the fact that a portion of PFC lies buried within the principal sulcus in this species and is inaccessible for laminar electrophysiology or optical imaging. The relatively lissencephalic cortex of the New World common marmoset (*Callithrix jacchus*) circumvents such limitations. It remains unknown however, whether marmoset PFC neurons exhibit persistent activity. Here, we addressed this gap by conducting wireless electrophysiological recordings in PFC of marmosets performing a delayed-match-to-location task on a home cage-based touchscreen system. As in macaques, marmoset PFC neurons exhibited sample, delay, and response-related activity that was directionally tuned and linked to correct task performance. Models constructed from population activity consistently and accurately predicted stimulus location throughout the delay period, supporting a framework of delay activity in which mnemonic representations are relatively stable in time. Taken together, our findings support the existence of common neural mechanisms underlying WM performance in PFC of macaques and marmosets, and thus validate the marmoset as a suitable model animal for investigating the microcircuitry underlying WM.

## Introduction

In 1936, Jacobsen tested Old World mangabey and baboon monkeys on a spatial delayed response task, in which the animal observed the hiding of food under one of two identical cups. After a delay of a few seconds to several minutes, the animal was allowed to choose one of the cups in order to obtain the food item. Jacobsen’s now-famous observation was that bilateral prefrontal cortex (PFC) lesions completely impaired the monkeys’ performance in this task. Subsequent studies narrowed the prefrontal regions responsible for maintenance of visuospatial information to the cortex surrounding the caudal two-thirds of the principal sulcus in rhesus macaque monkeys (Goldman and Rosvold 1970; Goldman et al. 1971). By recording from single neurons in this prefrontal region in macaque monkeys during similar manual spatial delayed response tasks, Fuster (1973), and Kubota and Niki (1971) found neurons that responded to the visual stimulus, anticipated the motor response, were active following the response, or, most surprisingly, neurons that discharged persistently during the delay period. In subsequent studies, Goldman-Rakic and colleagues took advantage of the behaviourally well-controlled oculomotor delayed-response task and demonstrated spatial tuning of delay-related activity and differences between correct and error trials (Funahashi et al. 1989, 1991). This led to the now-prevalent notion that persistent delay period activity in the dorsolateral PFC represents the cellular basis of spatial working memory (see for review Riley and Constantinidis 2016). Over the past 25 years, neural recordings in macaque monkeys have provided crucial insights into delay activity associated with various cognitive functions, and pharmacological studies in macaques have also begun to explore the influence of different neurotransmitter systems, such as dopamine (Ott and Nieder 2016) and acetylcholine (Vijayraghavan and Everling 2021), on persistent delay activity.

While macaque monkeys are an excellent animal model for studies of the neural basis of working memory in the PFC, the species also has several severe shortcomings: (1) a large part of the lateral PFC in macaques is deeply buried in the principal sulcus, making it difficult to access for laminar-specific recordings and manipulations, (2) macaques are large and difficult to handle, (3) they are expensive to house, (4) their low birth rate and long sexual maturation make longitudinal developmental studies challenging, and (5) pharmacological studies of new compounds are expensive due to the animal’s large body size. Although rodents, which also show delay-related activity in their frontal cortex, do not have these shortcomings, mice and rats lack a granular PFC. The so-called medial PFC of rodents likely corresponds to the primate anterior cingulate cortex and lacks the strong connectivity with parietal cortex that is characteristic of the primate lateral PFC. Moreover, it seems that delay-related activity in rodents is only present for short delays and is less robust than in macaque monkeys.

A nonhuman primate (NHP) species that avoids many of the problems of macaques, is the small New World common marmoset (*Callithrix jacchus*). The species is rapidly becoming popular as a powerful complement to canonical macaque and rodent models for preclinical modeling of the human brain in healthy and diseased states (Okano et al. 2016). The marmoset’s fast sexual maturation, low inter-birth interval, and routinely observed chimeric twinning make it the leading candidate for transgenic primate models (Sasaki et al. 2009; Okano et al. 2012; Kishi et al. 2014; Belmonte et al. 2015; Mitchell and Leopold 2015; Sasaki 2015). The lissencephalic (smooth) marmoset cortex also offers the opportunity for laminar electrophysiological recordings and optical imaging in frontoparietal areas. Most importantly, like macaques, marmosets have a granular lateral PFC that has strong connectivity with posterior parietal cortex (Burman et al. 2006; Reser et al. 2013).

Field studies have demonstrated that marmosets exhibit spatial working memory (Miles 1957; MacDonald et al. 1994; Tsujimoto and Sawaguchi 2002; Nakako et al. 2013) and touchscreen-based studies using delayed-match-to-location tasks (DML) have demonstrated that marmosets can maintain spatial information in working memory for at least several seconds (Spinelli et al. 2004, 2006; Yamazaki et al. 2016; Sadoun et al. 2019). In the DML task, a stimulus is briefly presented at one location of the screen (sample), followed by a waiting period (delay), and then the stimulus is again presented with one or several other identical stimuli (choice) and the subject has to touch the stimulus that is presented at the same location as the sample. The next step in establishing marmosets as an NHP model for PFC physiology is to test whether marmoset PFC neurons exhibit response profiles in delayed response task similar to those in macaques. To tackle this question, we trained three marmosets on the DML task using a custom-developed on-cage touchscreen system and wirelessly recorded neural activity using 96-channel Utah arrays chronically implanted over the marmoset PFC.

## Methods

### Subjects

Data was collected from three adult female common marmosets (*Callithrix jacchus*; Marmoset L, 41 months; Marmoset A, 26 months; Marmoset B, 24 months). All experimental procedures conducted were in accordance with the Canadian Council of Animal Care policy on the care and use of laboratory animals and a protocol approved by the Animal Care Committee of the University of Western Ontario Council on Animal Care. The animals were under the close supervision of university veterinarians.

### Delayed-Match-to-Location Task

Marmosets performed a delayed-match-to-location (DML) task (Figure 1) with varying delay lengths (2-8 s) on an in-house developed touchscreen testing box attached to the home cage (see *training protocol*, below). The touchscreen testing box measured 30 × 20 × 20 cm and had an opaque ceiling and floor. One end of the box was fitted with a metal docking plate designed to interface with the home cage and sliding door to allow access to the touchscreen testing box. Directly opposite to this was a slot for the touchscreen (254.80 × 177.50 mm, Elo 1002L). The remaining walls were constructed of transparent plexiglass which allowed light in and the marmosets to freely view the surrounding room space. For Marmosets A and B, we covered the box with a blanket as the room view appeared to cause stress and reduce their motivation to perform the task. A reward spout made from brass tubing was mounted on the floor with the outlet directly in front of the screen, 5-7 cm from the floor. As some marmosets preferred to use their snout to interact with the screen, to maintain consistency between marmosets we encouraged use of the forelimbs by affixing an array of vertical plexiglass bars (0.8 cm width, 2 cm spacing) in front of the screen. The task and reward delivery were controlled by a Raspberry PI 3 Model B running a custom-written Python script (marmtouch v0.0.1, Selvanayagam et al. 2021). Each trial began with the presentation of a sample stimulus (filled blue or pink circle, 3 cm diameter) on a grey background at one of four corner locations for 2.5 s (Figure 1). This was followed by a delay period in which the screen remained blank for 2-8 s. After the delay period, choice stimuli (filled blue or pink circles, 3 cm diameter) were presented at each of the four corner locations and the animal was required to touch the location matching the previously presented stimulus to obtain 0.075-0.1 mL of liquid reward per trial (50/50 mix of 1:1 acacia gum powder and water with liquid marshmallow). The reward was delivered via an infusion pump (model NE-510; New Era Pump Systems) through a liquid spout placed in front of the touchscreen monitor. Trials were separated by 5s intertrial periods. Animals performed 50-160 correct trials (lower and upper bound, respectively) per session for approximately 5-15 mL of liquid reward.

**Figure 1.**
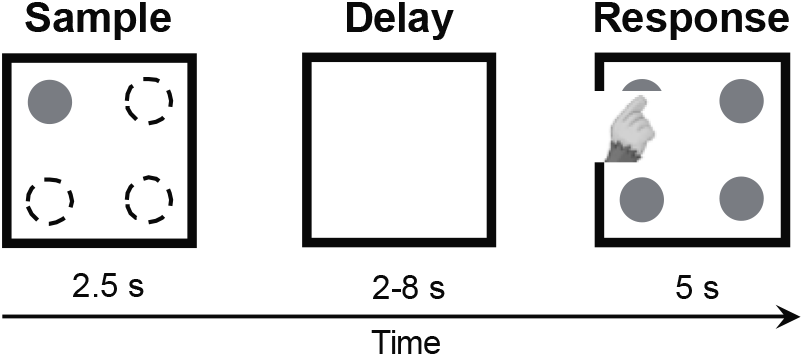
Delayed-match-to-location (DML) task. Successive panels indicate initial appearance of the sample stimulus, delay period and response (choice) phase. The sample was presented randomly in 1 of 4 possible locations.

### Training Protocol

Marmosets participated in a sequence of training phases derived via a process of successive approximations in order to establish their performance on the DML task. In a first training stage, we aimed to establish the association between touching a stimulus on the screen and obtaining a reward. To accomplish this, a small piece of marshmallow was placed at the centre of the screen, and a visual stimulus consisting of a large blue or pink circle (6 cm diameter) was presented at the same location. Touching the screen, and consequently the stimulus, allowed the animals to harvest the marshmallow and additionally triggered delivery of a liquid reward through the reward spout, which was paired with an auditory cue. Once they reliably performed this behaviour, we progressed to a stage in which the screen was no longer baited with a marshmallow, and touches to an identical large blue or pink circle were rewarded with liquid reward as in the previous stage. Because the DML task required the animals to touch stimuli at varying locations on the touchscreen, we next familiarized them with touching stimuli at varying locations. In this training phase, stimuli were presented randomly at one of the four corners of the touchscreen, and the animals received a liquid reward delivered from the reward spout for stimulus touches. Once this behaviour was performed reliably, we then introduced the concept of a short delay into the training protocol. In this phase, a visual sample stimulus was flashed at one location on the touchscreen. This was followed by a delay of 500 ms (incrementally increased to longer durations), after which a second comparison stimulus was flashed at the same location as the sample. Marmosets received a reward for touching this second comparison stimulus. To encourage exploratory responding and promote acquisition of the task concept, the animals were not restricted from touching the screen during the cue or delay periods. In addition, to simplify the task and training, we added the constraint that, in this stage and those following, cue and comparison stimuli were always presented at one of the four corners of the screen, rather than at locations randomized to any screen location on a trial-by-trial basis. We reasoned that this would make it easier for marmosets to acquire the task rule, while still requiring the use of mnemonic resources to correctly perform the task. Once the above steps were complete, animals advanced to the DML task proper, in which the sample and delay periods were identical to the previous training stage, but comparison stimuli were presented simultaneously at each of the four corners of the screen, forcing the animals to rely upon their mnemonic representations of the earlier sample location and knowledge of the location matching rule to perform the task correctly. However, an additional step was required for Marmoset A, as this animal remained at chance levels in the DML task proper. In this additional step, the animal was given a luminance cue to the correct location during the response period. A visual sample stimulus was presented at one location on the screen, followed by a delay during which the screen was blank. Following this, comparison stimuli were presented simultaneously at each of the four corners of the screen. Of these, the stimulus matching the earlier sample stimulus was presented at a slightly lower luminance, to act as an additional aid in acquiring the task rule of matching stimulus location. Once above chance levels, Marmoset A advanced to the DML task proper. Animals were able to complete this training protocol in 6-8 weeks.

### Array surgery

Once an animal reached 60 % correct task performance, it underwent an aseptic surgical procedure under general anesthesia in which a 96-channel Utah array (4 mm × 4 mm; 1 mm or 1.5 mm electrode length; 400 µm pitch; iridium oxide tips) were implanted in the left PFC (see Selvanayagam, 2019 for details). During this surgery, a microdrill was used to open a 4 mm burr hole in the skull and was enlarged as necessary using a rongeur. The dura was removed, and the array was manually inserted into the lateral PFC; wires and connectors were fixed to the skull using dental adhesive resin cement (All-Bond Universal and Duo-Link, Bisco Dental Products). Once implanted, the array site was covered with a thin layer of silicone adhesive (Kwik Sil; World Precision Instruments). A screw hole was drilled into the skull on the right side to place a stainless-steel ground screw. The ground wire of the array was then tightly wound around the base of the screw to ensure good electrical connection. A combination recording chamber/head holder (Johnston et al. 2018) was placed around the array and connectors and fixed in place using further layers of dental adhesive resin cement. Finally, a removable protective cap was placed on the chamber to protect the 3×32 channel omnetics connector.

### Neural recordings

After recovery from array implantation, we verified that electrode contacts were within the cortex by monitoring extracellular neural activity using the SpikeGadgets’ data acquisition system, Trodes (v2.2.2). Upon observing single-or multiunit activity at multiple sites of the array after about three weeks, we commenced unrestrained datalogger recordings of extracellular activity from the 96 implanted electrodes while the animal performed the DML task on a touchscreen attached to the home cage. Prior to a recording session, the animal was removed from the home cage. The protective chamber cap was removed, exposing the 3×32 omnetics connector. A custom-built routing board to three omnetics connector, horizontal headstage and an untethered datalogger (SpikeGadgets, San Francisco, US) was attached on top of the animal’s head. Once secured, the animal was returned to a smaller restricted area of the home cage where there was access to the touchscreen testing box for two hours. Subsequently, the animal was removed from the home cage for the removal of the routing board, headstage and datalogger. The protective chamber cap was placed on, and the animal was returned to the home cage. During the recording, the Logger Dock (SpikeGadgets, San Francisco, US) was used for untethered data acquisition and received synchronization pulses from the Raspberry Pi, aligned to sample onset. Camera(s) (Raspberry Pi camera (G) with fisheye lens or Arducam IMX477 Synchronized Stereo Camera with fisheye lens) were mounted on the in-house developed touchscreen testing box to record videos (aligned to sample onset) of the animal performing the DML task. In most recording sessions, animals were separated from their cage mate. However, in a few sessions where the animals were not separated (to reduce stress from separation), recorded videos were used to score each trial for animal identity.

Neural data were first filtered with a common median filter to remove large movement related artifacts. This was then filtered with a 4-pole Butterworth 500 Hz high-pass filter. Spike detection and sorting was performed offline (Plexon Offline Sorter v3). Only clearly isolated single units with baseline discharge rates greater than 0.5 Hz were included in the analysis (149/1853 units were excluded). Videos of all trials in all sessions were manually scored to determine whether the animal looked at the sample during the sample-presentation period. Only experimental sessions with at least 10 correct trials, where the animal looked at the sample during the sample-presentation period were included (5 of 33 sessions were excluded).

### Data analysis

Analysis was performed using custom code written in Matlab (MathWorks).

Activity within distinct task epochs was assessed using analyses of variance (ANOVA) to compare the mean discharge rates of each neuron for each condition (four spatial locations) and each task epoch (baseline, sample, delay, pre-response and post-response). The baseline epoch was defined as the 1.5 s prior to the onset of the sample stimulus to the time of sample stimulus presentation. The sample epoch was defined as the 100 ms after the onset of the sample stimulus to the time of stimulus offset (2.4 s). We excluded the first 100 ms to avoid any potential sluggish sample-related activity contaminating estimates of delay-related activity. The delay epoch was 4 s for Marmoset L and 2 s for Marmoset A and B. The pre-response epoch was defined as the 300 ms period prior to the touch response and the post-response epoch was defined as the 1000 ms immediately after the touch response. Statistical significance was evaluated at an alpha level of p < .02.

To investigate differences in activity between correct and error trials, a modulation index was computed by subtracting the discharge rate of the non-preferred condition from the discharge rate of the preferred condition, divided by the sum of the discharge rate of the preferred and non-preferred condition. For each unit, we classified the preferred and non-preferred conditions as those in which the unit had the highest and lowest mean discharge rate during the delay epoch in the correct trials respectively.

### Waveform preprocessing and cell type classification

For each single unit, the mean waveform was interpolated (cubic spline) from an original sampling rate of 30 kHz to 1 MHz. For cell class classification, we computed two measures of the resultant waveform: trough-to-peak duration and time for repolarization. The time for repolarization was defined as the time at which the waveform amplitude decayed 25% from its peak value (Ardid et al. 2015). To identify broad spiking (BS) and narrow spiking (NS) cell clusters in the data we used the unsupervised algorithm used by Trainito and colleagues (2019).

In this method, the expectation-maximization (EM) algorithm is used to estimate the parameters of the gaussian mixture model (GMM), a statistical model which describes the mean and variance of underlying subpopulations.

### Pattern Classifier

To determine whether delay activity was sustained throughout the entire delay period, for each session we trained linear supply vector machines (SVM) to predict the sample stimulus location using the discharge activity of each neuron in a sliding window over the task interval. We constructed models for 1 s time bins starting 1.5 s before sample onset (baseline) to 1 s after the end of the delay epoch in steps of 500 ms. Models were trained with 80 % of the data at a specific time bin and tested at the same time bin with the remaining 20 % of data for a classification accuracy. The same model was used to test the same 20 % of data at all other time bins. This process of separating the data into a training and testing set was repeated 100 times and the classification accuracies were averaged to provide a more robust estimate. These values were then averaged across sessions for each animal separately.

To investigate differences in the two populations of neurons (BS vs NS), we trained a single model over the entire delay period using the entire dataset for each session. Note that since we used a one vs one linear SVM, each feature (neuron) has a coefficient for each pair of conditions (4 choose 2, 6 coefficients). As a measure of each unit’s contribution to the model, we computed the magnitude of the 6-dimensional vector constructed from these coefficients. A larger magnitude suggests that a neuron’s activity is better able to separate between all pairs of conditions. We then compared these weights between BS and NS cells in an independent samples t-test.

### Array localization

Marmosets were euthanized at the end of the data acquisition process to prepare the brains for ex-vivo MRI scan. The animals were deeply anesthetized with 20 mg/kg of ketamine plus 0.025 mg/kg medetomidine and 5% isoflurane in 1.4–2% oxygen to reach beyond the surgical plane (i.e., no response to toe pinching or cornea touching). They were then transcardially perfused with 0.9% sodium chloride irrigation solution, followed by 4% paraformaldehyde in 0.1 M phosphate buffer solution or 10% buffered formalin. The brain was then extracted and stored in 10% buffered formalin for more than a week before ex-vivo magnetic resonance imaging (MRI). On the day of the scan, the brain was transferred to another container for imaging and immersed in a fluorine-based lubricant (Christo-lube; Lubrication Technology) to improve homogeneity and avoid susceptibility artifacts at the boundaries. Ex-vivo MRI was performed on a 9.4T 31 cm horizontal bore magnet (Varian/Agilent, Yarnton, UK) and Bruker BioSpec Avance III console with the software package Paravision-7 (Bruker BioSpin Corp, Billerica, MA), a custom-built high-performance 15-cm-diameter gradient coil with 400 mT/m maximum gradient strength (xMR, London, CAN; Peterson et al. 2018), and an mp30 (Varian Inc., Palo Alto, USA) transmit/receive coil. High resolution (100×100×100 µm for Marmoset A and Marmoset B, 100×100×200 µm for Marmoset L) T2-weighted images were acquired for each animal. The raw MRI images were converted to NifTI format using dcm2niix (Li et al. 2016) and the MRIs were nonlinearly registered to the ultra-high-resolution ex-vivo NIH template brain (Liu et al. 2018), that contains the location of cytoarchitectonic boundaries of the Paxinos atlas (Paxinos et al. 2012), using Advanced Normalization Tools (ANTs; Avants et al. 2011) software. The resultant transformation matrices were then applied to the cytoarchitectonic boundary image included with the NIH template brain atlas. These cytoarchitectonic boundaries overlayed on the registered ex-vivo anatomical T2 images were used to reconstruct the location of the implanted array in each marmoset (Supplementary Figure 1).

## Results

### Task performance

Marmosets performed a total of 28 sessions (Marmoset L, 18 sessions; Marmoset B, 8 sessions; Marmoset A, 2 sessions). Overall, we observed that marmosets were able to perform the DML task at above chance accuracy at a range of delay durations. Data were collected from three animals at a 2s delay, (Marmoset L = 70.6 %, Marmoset B = 64.1 %, Marmoset A = 64.2 %). Marmoset L additionally performed the task at two longer delay durations (4 s and 8 s). Consistent with many reports, performance declined with increasing durations, falling to 63.4 % and 54.2 %, respectively. This observation was confirmed statistically via one-way ANOVA (F(2,23) = 12.49, *p* < .001). A *post hoc* Bonferroni corrected t-test revealed that mean task accuracy was significantly lower following 8s delays as compared to 2 s or 4 s delays (*p* < .05).

To further investigate task performance, we compared marmosets’ reaction times (RT) on correct and error trials at the 2 s delay duration. The logic of the DML task, as in other delayed-response tasks, is that animals acquire a mnemonic representation of the spatial location of the cue stimulus during cue presentation, maintain this representation during the delay period of the task, and subsequently compare this representation with the stimuli presented during the response period to select the correct response. Generally speaking, if animals are relying on mnemonic processes to guide response selection, it is expected that reaction times on error trials will be similar to or longer than those on correct trials, since they should reflect “diligent guesses” regarding the correct location (Link 1982). For the 2s delay condition we found that in all cases, RTs on error trials were equal to or greater than those on correct trials, with RT’s significantly greater for error than correct trials for Marmosets L and A (t-test, p < .05) (Marmoset L: x□ correct RT = 1.23s, x□ error RT = 1.99s, Marmoset B: x□ correct RT = 1.43s, x□ error RT = 1.57s, Marmoset A: x□ correct RT = 1.20s, x□ error RT = 1.84s). These results are consistent with a reliance of marmosets on mnemonic processes during DML task performance. Taken together with the findings of above chance task performance for all animals, and the decline in task performance as a function of delay duration observed in Marmoset L, these data indicate that performance on the DML task was an accurate reflection of Marmosets’ spatial WM abilities.

### Marmoset PFC Neurons Exhibit Sample, Delay, and Response Period Activity

To determine whether marmoset PFC neurons exhibit task-related activity in the DML task, we simultaneously recorded the activity of a total of 1704 PFC neurons (502 in Marmoset L, 908 in Marmoset B, 294 in Marmoset A) in three monkeys over 28 experimental sessions in which they performed this task. We included in our analyses only well-isolated single units with baseline discharge rates greater than 0.5 Hz. Many neurons were modulated during the DML task (460 in Marmoset L, 782 in Marmoset B, 209 in Marmoset A), which we defined as a significant difference in mean discharge rate between a task epoch and a baseline period. Representative single neurons exhibiting modulations during the sample epoch, delay epoch, both sample and delay epoch, as well as during the response epochs are presented in Figures 2 and 3. Many neurons exhibited task-related activity modulations, and these modulations were observed in all epochs of the DML task, with a bias toward units responding in the two response epochs. Of the 460 neurons modulated during the task in Marmoset L, 135 neurons (26.9 % of overall neurons) displayed sample-related activity, 238 neurons (47.4 %) displayed delay period activity, 359 neurons (71.5 %) displayed pre-response related activity and 402 neurons (80.1 %) displayed post-response related activity. In Marmoset B, 247 (27.2 %), 228 (25.1 %), 557 (61.3 %) and 630 (69.4 %) neurons, as well as in Marmoset A, 83 (28.2 %), 67 (22.8 %), 130 (44.2 %) and 159 (54.1 %) neurons displayed sample-related activity, delay period activity, pre-response related activity and post-response related activity, respectively. Overall, we found that, across the three animals, well-isolated single units were recorded across a broad set of prefrontal subregions including areas 46D, 46V, 8aD, 8aV, 9, and 10 (see Figure 4). We observed task-related activity in all epochs in all subregions but noted that the proportion of units with delay activity was relatively lower in areas 9 and 10.

**Figure 2.**
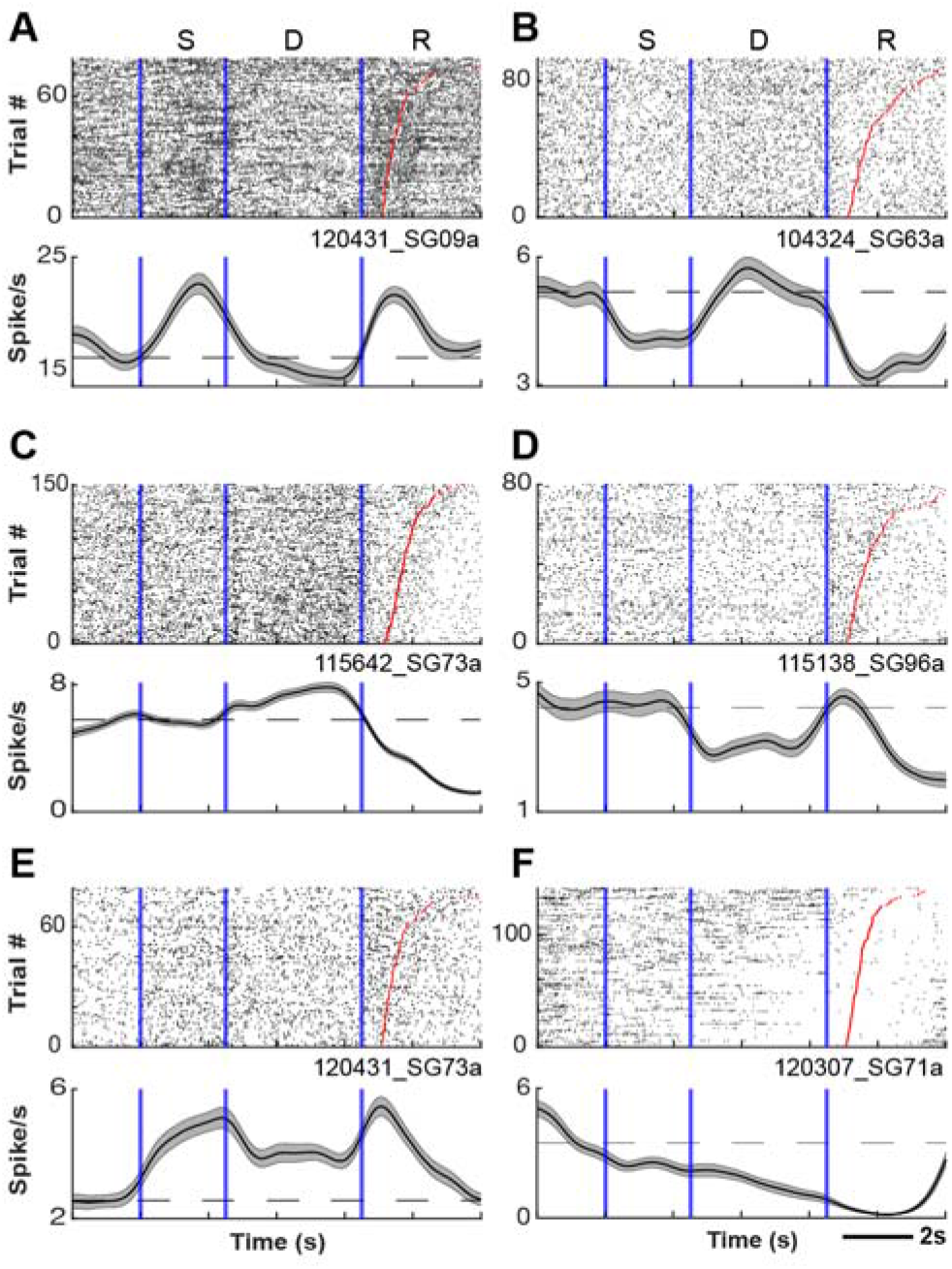
PFC neurons exhibit sample- and delay-period activity during the DML task. Representative single neurons exhibiting excited**(A)** or suppressed **(B)** activity during the sample epoch, excited **(C)** or suppressed **(D)** activity during the delay epoch, excited **(E)** or suppressed **(F)** activity during both sample and delay epochs. Rasters are aligned to sample-onset. Red lines depict reaction time.

**Figure 3.**
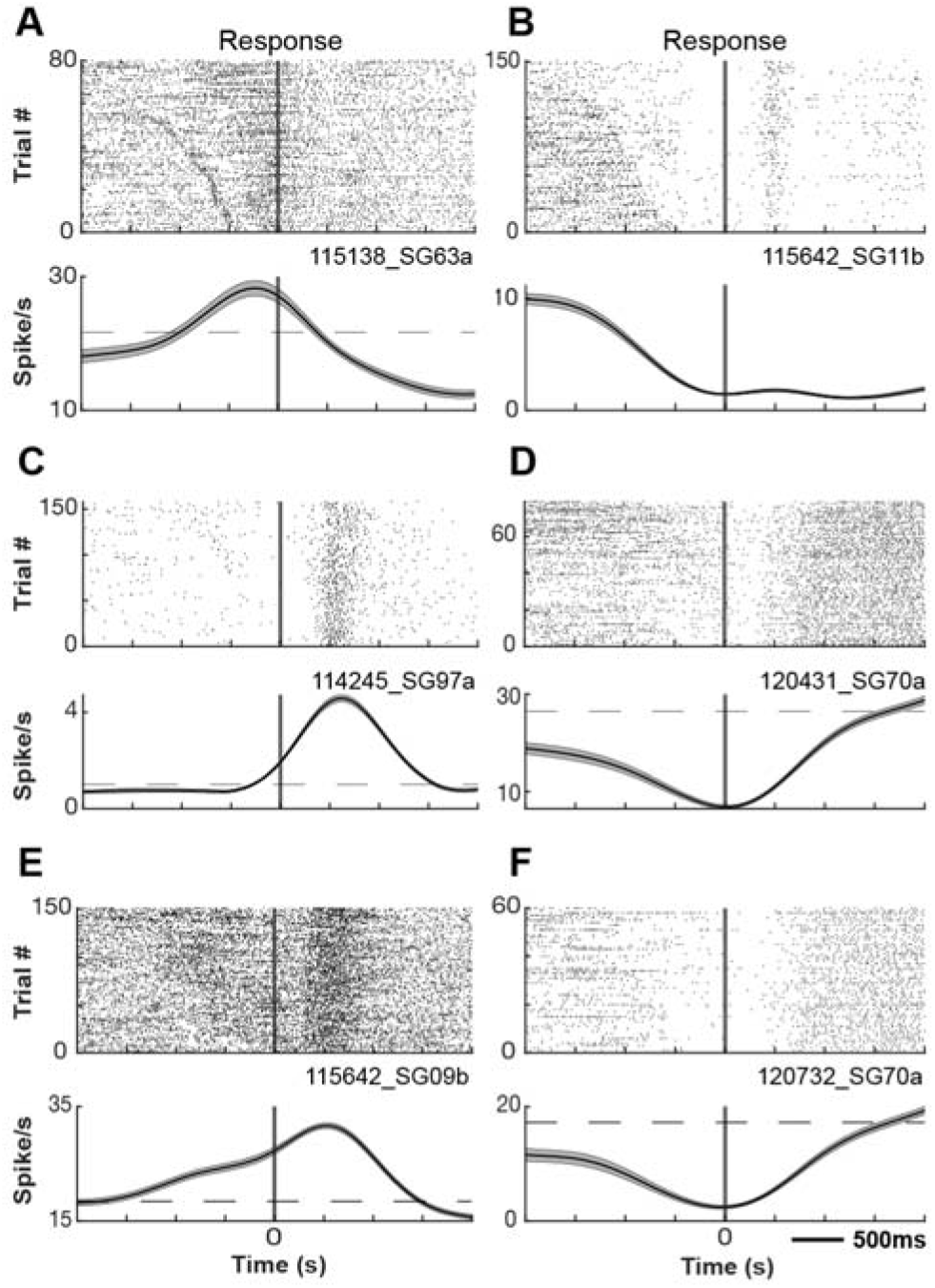
PFC neurons exhibit response-related activity during the DML task. Representative single neurons exhibiting excited **(A)** or suppressed **(B)** activity during the pre-response epoch, excited **(C)** or suppressed **(D)** activity during the post-response epoch, excited **(E)** or suppressed **(F)** activity during both pre- and post-response epochs. Rasters are aligned to response.

**Figure 4.**
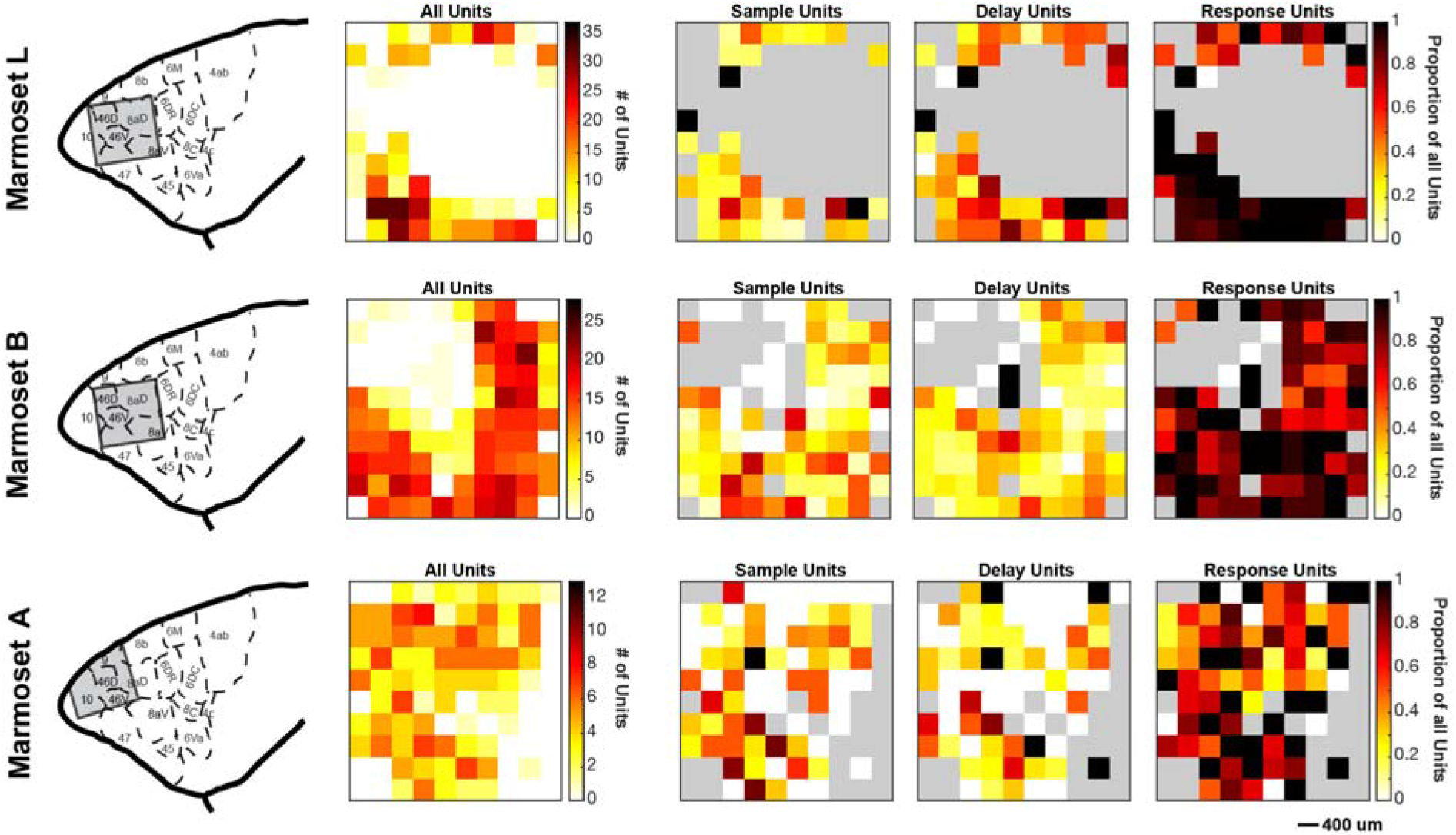
Distribution of task-modulated units across recording arrays. Array locations were reconstructed using high-resolution MRIs and superimposed on a standardized marmoset brain, area boundaries from Paxinos et al. (2012; left panel). For each marmoset, task modulated units were distributed across the array in locations where well-isolated single units were observed, followed by locations on the array where sample-, delay- and response-related units were observed (from left to right). Grey colour depicts where on the array well-isolated single units were not observed.

Prior work in macaque monkeys performing delayed-response tasks has revealed that modulations of sample-, delay-, and response-related activity may take the form of either increases or decreases from the baseline discharge rate (Funahashi et al. 1989, 1991). To investigate such modulations in marmoset PFC, we further classified whether each neuron was significantly excited or suppressed in each task epoch with respect to baseline (Table 1). The proportions of neurons excited and suppressed significantly varied across task epochs (chi-square test, Marmoset L: *X*^2^ (3, *N* = 1134) = 37.49, *p* < .001; Marmoset B: *X*^2^ (3, *N* = 1662) = 19.41, *p* < .001; Marmoset A: *X*^2^ (3, *N* = 439) = 5.57, *p* < .001). Overall, these data show that marmoset PFC neurons are modulated in a similar manner to those in macaque during the DML task.

**Table 1.** Number of neurons excited or suppressed in each task epoch for each marmoset.

| Marmoset | Modulation | Epoch |  |  |  |
| --- | --- | --- | --- | --- | --- |
|  |  | Sample | Delay | Pre-response | Post-response |
| L | Excited | 63 (47%) | 95 (40%) | 91 (25%) | 99 (25%) |
|  | Suppressed | 72 (53%) | 143 (60%) | 268 (75%) | 303 (75%) |
| B | Excited | 97 (39%) | 49 (221%) | 171 (31%) | 176 (28%) |
|  | Suppressed | 150 (61%) | 179 (79%) | 386 (69%) | 454 (72%) |
| A | Excited | 36 (43%) | 19 (28%) | 39 (30%) | 49 (31%) |
|  | Suppressed | 47 (57%) | 48 (72%) | 91 (70%) | 110 (69%) |

### Marmoset PFC Neurons Exhibit Spatial Tuning During DML Task Performance

A now-classic observation in macaque PFC is that single units exhibit spatial tuning of discharge rates in the sample, delay, and response epochs of spatial working memory tasks. We similarly observed such tuning in marmoset PFC neurons in all animals from which we recorded. To investigate this statistically, we carried out separate one-way ANOVAs with sample location at 4 levels for each task epoch separately. An example neuron exhibiting such tuning is depicted in Figure 5. Overall, in Marmoset L, 32 neurons (23.7 %, proportion in respect to the number of neurons modulated in the epoch) displayed tuning during the sample period, 58 neurons (24.4 %) displayed tuning during the delay period, 80 neurons (22.3 %) displayed tuning during the pre-response period and 135 neurons (33.6 %) displayed tuning during the post-response period. In Marmoset B. 45 (18.2 %), 20 (8.8 %), 143 (25.7 %) and 204 (32.4 %) neurons, as well as in Marmoset A, 2 (2.4 %), 10 (14.9 %), 16 (12.3 %) and 18 (11.3 %) neurons displayed tuning activity during the sample period, delay period, pre-response period and post-response period, respectively.

**Figure 5.**
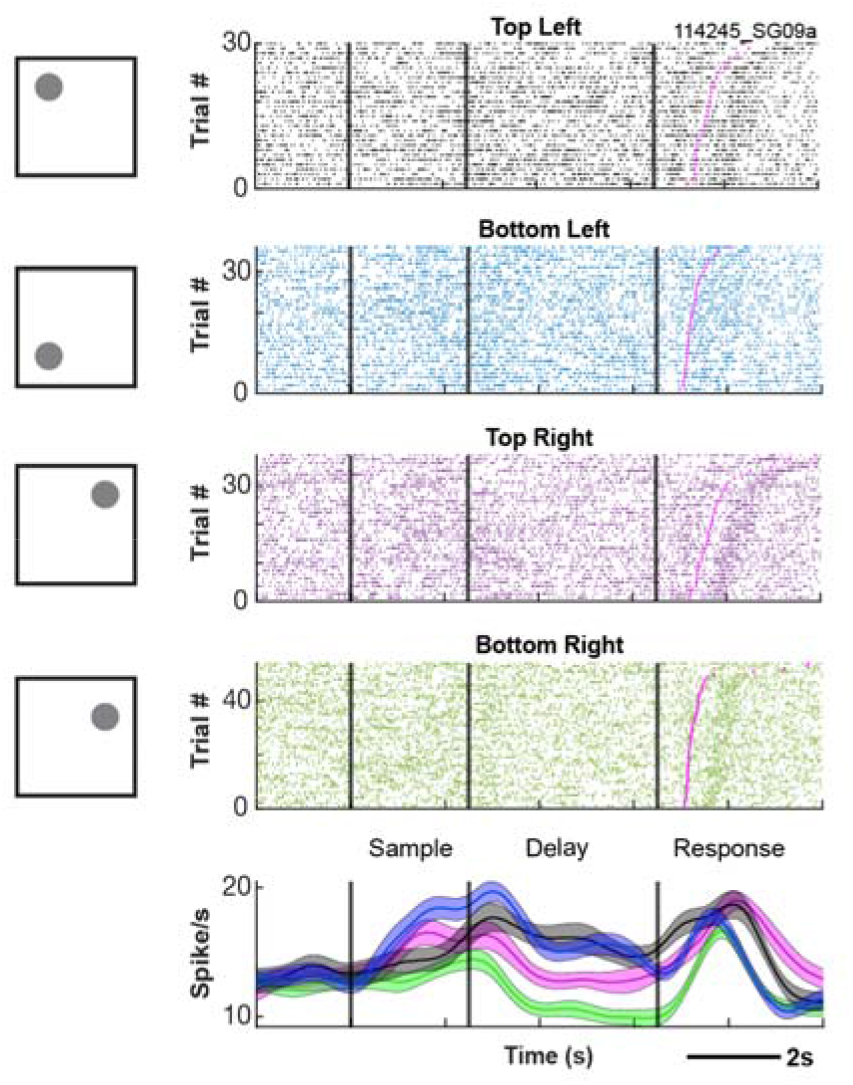
Example neuron exhibiting spatial tuning during the DML task. Rasters are aligned to sample-onset.

### Neural activity reflects mnemonic processes rather than simple orienting

One possibility, noted also in earlier work (see Goldman-Rakic 1995), was that animals were able to correctly perform the task not by relying on WM *per se*, but rather by simply orienting the head or whole body toward the location of the stimulus during sample presentation, and maintaining that orientation throughout the delay period until responding. This strategy would not only circumvent the cognitive process of WM, but additionally potentially contaminate any observations of delay period activity with a sustained signal indicating body or head orientation. To address this, in several sessions (3 in Marmoset L, 2 in Marmoset B, 2 in Marmoset A), videos of all correct trials were manually scored with respect to whether the animal was orienting (head-oriented) towards the sample during the delay-period (Supplementary Video 1). or not (Supplementary Video 2). In the majority of trials (95.8 % for Marmoset L, 92.3 % for Marmoset B, 94.9 % for Marmoset A), the animals did not fixate on the location of the sample during the delay-period. We removed the trials on which the animals fixated on the location of the sample and performed an ANOVA on condition and task epoch again to compare the differences. For these sessions, the number of neurons that displayed delay-period activity was similar (Marmoset L: 55/59 neurons (93.2 %); Marmoset B: 81/85 neurons (95.3 %); Marmoset A: 62/67 neurons (92.5 %), indicating that our observations were not contaminated by orienting or strategic processes and reflect instead the retention of mnemonic information.

### Persistent Delay Period Activity in Marmoset PFC Reflects Task Performance

A seminal observation linking persistent activity and task performance is that persistent delay period activity is attenuated on trials on which animals make performance errors relative to those on which they perform correctly (Funahashi et al. 1989). To investigate this in marmoset PFC, we compared discharge rates between correct and error trials for neurons which were responsive during the delay period (e.g. exhibited discharge rates significantly elevated during the delay epoch relative to baseline). In Marmoset L, 119/204 neurons (58.3 %) exhibited a significant difference in discharge rate between correct and incorrect trials during the delay period. In Marmoset B, 48/212 (22.6 %) and in Marmoset A, 28/63 (44.4 %) neurons had a significant difference (independent samples t-test, p <.05). Figure 6A depicts an example neuron for which the discharge rate during the delay period was significantly different between correct and incorrect trials.

**Figure 6.**
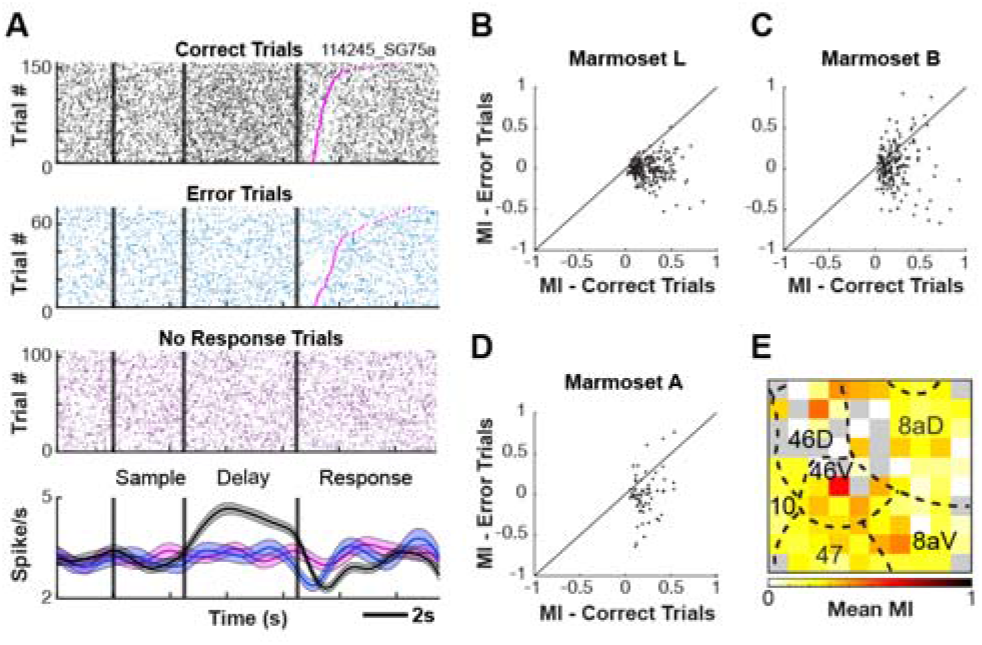
Delay-period activity is reduced on error trials. Example neuron for which the discharge rate during the delay-period is significantly different between correct and error trials **(A)**. Delay-period modulation index computed from the preferred and non-preferred conditions separately for correct and error trials **(B-D)**. Average modulation indices of neurons recorded at each electrode contact of the Utah array across sessions for Marmoset L and A **(E)**. Grey colour depicts where on the array well-isolated single units were not observed.

**Figure 7.**
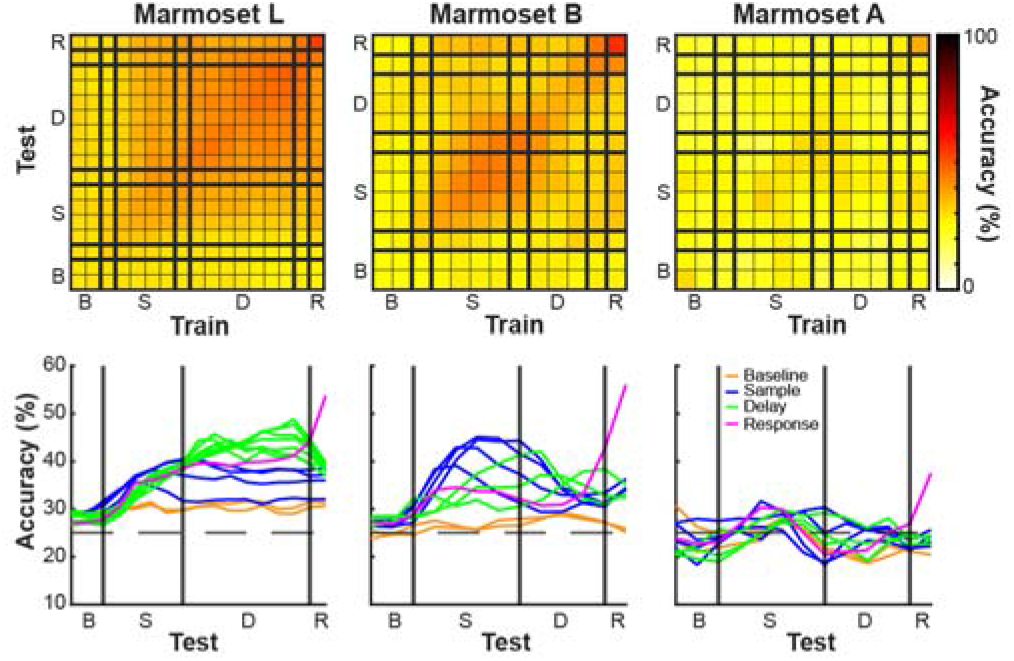
Stable versus dynamic mnemonic representations at the population level. For each session we trained a pattern classifier to predict the sample stimulus location from the population activity in a sliding window over the task interval (trained on data from a specific time point and tested on the same and different time points). Above chance classification accuracy (25 %) indicates that a representation of the information being classified exists in the population. Stable population coding was present during the delay period in Marmoset L and B, but not in Marmoset A.

To further investigate the magnitude of differences in neural activity between correct and incorrect trials during the delay period we computed a modulation index from the preferred and non-preferred conditions separately for correct and incorrect trials (Figure 6B). For Marmoset L, the modulation index was higher for correct trials than incorrect trials (0.2414 vs -0.0065, *p* < .001). This trend was similar for Marmoset B (0.222 vs 0.034, *p* < .001) and Marmoset A (0.230 vs -0.018, *p* < .001). Overall, these data are consistent with previous reports indicating that delay period activity is reduced on error trials, and provide evidence for a link between the magnitude of persistent delay period activity and WM task performance.

To determine the PFC locations at which we observed the strongest, spatially selective delay activity, we averaged the modulation indices of neurons recorded at each electrode contact of the Utah array across sessions. We pooled across the neurons recorded from Marmosets L and B to improve our sampling of common PFC subregions as the arrays in these animals shared a high degree of overlap with respect to the PFC areas sampled (See Figure 4). We observed the greatest mean modulation indices at the center of the array along the anterior-posterior axis, roughly corresponding to areas 46, 8aV and 8aD (see Figure 6E).

### Broad and Narrow Spiking Neurons are Modulated in all Task Epochs

To investigate the contributions of putative pyramidal cells and interneurons to mnemonic processes in marmoset PFC, we used the expectation-maximization algorithm for Gaussian mixture model clustering method on the trough-to-peak duration and time for repolarization of single unit waveforms to identify BS and NS cell clusters in an unsupervised manner (Supplementary Figure 2). This resulted in 414 cells (28.5 %) being classified as narrow spiking and 1037 cells (71.5 %) being classified as broad spiking cells (Marmoset L: BS = 286 cells, NS = 174 cells; Marmoset B: BS = 575 cells, NS = 207 cells; Marmoset A: BS = 176 cells, NS = 33 cells). For neurons modulated in each epoch of the DML task, we identified whether the neuron was classified as BS or NS. In Marmoset L, 94 BS cells (69.6 %) displayed sample-related activity, 153 BS cells (64.3 %) displayed delay period activity, 223 BS cells (62.1 %) displayed pre-response related activity and 254 BS cells (63.2 %) displayed post-response related activity. In Marmoset B, 180 (72.9 %), 162 (71.1 %), 407 (73.1 %) and 466 BS cells (74.0 %), as well as in Marmoset A, 67 (80.7 %), 59 (88.1 %), 108 (83.1 %) and 132 BS cells (83.0 %) displayed sample-related activity, delay period activity, pre-response related activity and post-response related activity, respectively. For the task modulated cells, Chi-Square Goodness of Fit Tests were conducted to determine if the proportion of each cell type varied across the task epochs. No significant differences were observed (Marmoset L: *X*^2^ (3, *N* = 1134) = 2.52, *p* > .05; Marmoset B: *X*^2^ (3, *N* = 1662) = 0.73, *p* > .05; Marmoset A: *X*^2^ (3, *N* = 439) = 1.50, *p* > .05). We further identified whether BS and NS cells were excited or suppressed in each task epoch compared to baseline (Table 2). Overall, consistent with previous reports in macaque PFC (Wilson et al. 1994; Rao et al. 1999; Constantinidis et al. 2002), we observed that both BS and NS neurons were modulated in all task epochs, and that this modulation could take the form of excitation or suppression.

**Table 2.** Number of neurons excited or suppressed in each task epoch for each marmoset separated by cell type.

| Marmoset | Modulation | Cell Type | Epoch |  |  |  |
| --- | --- | --- | --- | --- | --- | --- |
|  |  |  | Sample | Delay | Pre-response | Post-response |
| L | Excited | BS | 34 | 58 | 58 | 62 |
|  |  | NS | 29 | 37 | 33 | 37 |
|  | Suppressed | BS | 60 | 95 | 165 | 192 |
|  |  | NS | 12 | 48 | 103 | 111 |
| B | Excited | BS | 79 | 45 | 118 | 132 |
|  |  | NS | 18 | 4 | 53 | 44 |
|  | Suppressed | BS | 101 | 117 | 289 | 334 |
|  |  | NS | 49 | 62 | 97 | 120 |
| A | Excited | BS | 29 | 17 | 31 | 40 |
|  |  | NS | 7 | 2 | 8 | 9 |
|  | Suppressed | BS | 38 | 42 | 77 | 92 |
|  |  | NS | 9 | 6 | 14 | 18 |

### Stable vs Dynamic Mnemonic representations in Persistent Activity

To determine whether delay activity was stable in time or temporally dynamic, for each session we trained a pattern classifier to predict the sample stimulus location from the population activity in a sliding window over the task interval. The classifier can be trained on data from a specific time point in the trial and can be tested on the same time point, or different time points, for a measure of the stability of the representation over the course of WM maintenance (Sreenivasan and D’Esposito 2019) . Above chance classification accuracy (25 %) indicates that a representation of the information being classified exists in the population. In Marmosets L and B, classifiers trained at a given delay epoch time bin predicted stimulus location in other delay epoch time bins more accurately than chance; this was not the case for Marmoset A (see Figure 9). The results indicate that stable population coding was present during the delay period in Marmoset L and Marmoset B, but not in Marmoset A.

We separated the population of neurons within each session into two categories (neurons that display delay-period activity and neurons that did not as determined above) to determine their influence on the trained model. As we had used a one versus one, linear SVM with four classes (stimulus location) where each feature (neuron) had 6 (4 choose 2) coefficients, to determine a measure of each unit’s contribution to the model, we computed the magnitude of the 6-dimensional vector constructed from these coefficients. These weights were greater for neurons that displayed delay-period activity (Marmoset L: Mean = 1.67, SE = 0.06; Marmoset B: Mean = 0.98, SE = 0.03) as compared to neurons that did not display delay-period activity (Marmoset L: Mean = 1.43, SE = 0.04; Marmoset B: Mean = 0.90, SE = 0.02; t-test: Marmoset L: p < .001; Marmoset B: p < .05). The results indicated that neurons that display delay-period activity are mostly responsible for the stability of the representation over the course of WM maintenance.

We additionally separated the population of neurons within each session into cell types and performed the same analysis during the delay-period. These weights were greater for broad spiking cells (Marmoset L: Mean = 1.62, SE = 0.04; Marmoset B: Mean = 0.97, SE = 0.02) as compared to narrow spiking cells (Marmoset L: Mean = 1.40, SE = 0.05; Marmoset B: Mean = 0.79, SE = 0.02; t-test: Marmoset L: p < .01; Marmoset B: p < .001). The results suggested that broad spiking cells contributed more than narrow spiking cells to the population’s representation of stimulus location throughout the delay-period.

## Discussion

Activity during delay periods is a hallmark of macaque prefrontal cortex (Fuster and Alexander 1971; Fuster 1973; Niki and Watanabe 1976; Quintana et al. 1988; Funahashi et al. 1989, 1990). Although delay-related activity has been reported in rodents, it is only present for short delay periods and it is substantially less robust than in macaques. Whether small New-world common marmosets show delay-related activity comparable to Old-world macaques remains unanswered. Here we recorded single neuron activity in the marmoset prefrontal cortex while unrestrained monkeys performed a touchscreen-based delayed-match-to-location (DML) task to address this question. We found that common marmosets possess robust delay-related activity throughout their prefrontal cortex and that this activity seems to be sustained throughout delay periods.

The first delay-related activity in macaques was recorded in head-restrained monkeys performing a manual spatial delayed response task (Fuster and Alexander 1971; Kubota and Niki 1971). Subsequently, an eye movement version of the spatial delayed response task, the oculomotor delayed response (ODR) task was popularized by Goldman-Rakic’s lab (Funahashi et al. 1989, 1990, 1991). The ODR task has the distinct advantage over previous manual versions in that eye positions can be controlled and samples can be presented and maintained in known retinotopic coordinates (Goldman-Rakic 1995). Although marmosets can be trained to perform saccadic eye movement tasks, it has thus far proven difficult to train these monkeys on the ODR task. To date, there are only a few conference reports regarding training marmosets on the ODR task, and the consensus is the performance of marmosets is quite low and that delay length is limited to short periods under 400 ms (Amly et al. 2021). Another group (Carney et al. 2019) delivered reward throughout the sample and delay periods as a means to encourage marmosets to maintain fixation, which often presents a challenge for this primate species and detrimentally affects task performance. In that case they reported performance comparable to macaques, although the delivery of rewards concurrent with the delay period of task potentially confounds the interpretation of persistent delay activity since reward-related and mnemonic signals would be intermixed. We have also not succeeded in training marmosets on the ODR task. In contrast to the difficulties inherent in employing the ODR task in marmosets, several studies have shown that marmosets can be trained on the touchscreen version of a DML task with delay periods of up to 68s (Spinelli et al. 2004, 2006; Yamazaki et al. 2016; Sadoun et al. 2019).

The delay-related activity that we observed in marmoset PFC neurons is remarkably similar to the initial reports by Fuster and Kubota (Fuster and Alexander 1971; Kubota and Niki 1971) in the macaque. Like macaque PFC neurons, marmoset PFC neurons exhibit sample-, delay-, and response-related activity. The profiles of individual neurons closely resembled those described in macaques. Some were active in the sample and response period, some were active during the sample and delay period, and other were active just during the delay period (Figure 2) While we found many neurons that increased their activity during these periods, we also found an even larger proportion of neurons that decreased their activity from baseline during the different task epochs. This is also very similar to reports in macaques.

Due to the small size of the PFC in marmosets, each of our implanted 4×4 mm arrays covered multiple prefrontal areas. This allowed us to compare the density of task-related activity during different epochs across different areas. Although we found delay-related activity in all three animals in all sampled areas, including area 8Ad, 8Av, 47, 46V, 46D, and 10, it was considerably weaker in marmoset A in which the array was implanted further anterior and medial than in the other two animals. This suggests that delay activity is less robust in marmoset areas 9 and 10. The strongest sample- and delay-related activity was found in areas 46, 47 and 8Av. Parts of area 8Av correspond to the frontal eye fields (FEF) in the marmoset (Selvanayagam et al. 2019). In fact, Marmoset L was also one of the subjects in our previous electrical microstimulation study and fixed vector saccades could be evoked at posterior lateral electrodes of this array. This is consistent with findings in macaques which also show robust delay-related activity at sites corresponding to the FEF in area 8 (Funahashi et al. 1989).

A distinguishing feature of delay cells in primate lateral prefrontal cortex is spatial tuning for the cued location (Funahashi et al. 1989, 1990; Funahashi 2006). Here, we observed spatial tuning in many of our delay cells and this effect was significantly stronger on correct than on error trials supporting the hypothesis that delay-related activity is linked to task performance in marmosets. Additionally, the large number of electrodes on our Utah arrays allowed us to sample a large number of cells simultaneously facilitating population level analyses. We were able to predict stimulus location from the population activity above chance levels, demonstrating for the first time that delay-related activity in marmoset prefrontal cortex represents mnemonic information at the population and single neuron levels.

Since the first recording studies by Fuster and Kubota (Fuster and Alexander 1971; Kubota and Niki 1971), working memory has been thought to be mediated by persistent sustained neuronal spiking during the delay period (Goldman-Rakic 1995), and many circuit models of recurrent prefrontal circuits have been developed to understand the mechanisms underlying such activity (see Wang 2021). An alternative proposal is that working memory is instantiated by discrete spiking bursts rather than sustained activity (Lundqvist et al. 2016). Here our results suggest that persistent activity was sustained throughout the delay period supporting a framework of delay activity in which mnemonic representations remain relatively stable in time.

We additionally observed that broad spiking neurons contributed more to the population’s representation of the sample stimulus’ spatial location than narrow spiking neurons. This finding dovetails with previous reports in macaques performing the ODR task demonstrating broader tuning of narrow than broad spiking neurons, and substantially higher noise correlations between narrow spiking putative interneurons than broad spiking putative pyramidal neurons (Constantinidis and Goldman-Rakic 2002). This is also consistent with the notion that the presence of correlated noise reduced the representation of spatial location in narrow relative to broad spiking neurons (Averbeck and Lee 2006).

In summary, we have shown that neural activity in the marmoset PFC can be chronically recorded in touchscreen tasks using completely unrestrained datalogger-based recording technology. In this first study, our goal was to characterize the activity profiles of PFC neurons in marmosets and to determine the distribution of sample, delay, and response-related activity across prefrontal regions. Subsequent studies can exploit the lissencephalic marmoset PFC to employ laminar electrophysiological (Jun et al. 2017; Johnston et al. 2019) or miniscope recordings using implanted prism lenses (Kondo et al. 2018) in touchscreen tasks to characterize the functional microcircuitry during working memory tasks in different PFC regions in unrestrained marmosets. Combined experiments with optogenetic manipulations promise to provide further insights into the microcircuit mechanisms of delay-related activity in the primate PFC.

## Acknowledgements

The authors wish to thank Cheryl Vander Tuin, Whitney Froese, and Hannah Pettypiece for animal preparation and care and Peter Zeman for technical assistance. We would also like to thank David Everling for assistance with touchscreen testing. This research was supported by the Canadian Institutes of Health Research (CIHR) grant FRN148365 to SE and the Canada First Research Excellence Fund to BrainsCAN. RW was also supported by the Canada First Research Excellence Fund to BrainsCAN and the Next Generation Networks for Neuroscience (NeuroNex). JS was supported by a Natural Sciences and Engineering Research Council (NSERC) Canadian Graduate Scholarship (Doctoral).

## References

Amly W, Chen C-Y, Onoe H, Isa T. 2021. Dissecting errors of the exogenously-driven and endogenously-driven saccadic tasks in the common marmosets (preprint). Neuroscience.

Ardid S, Vinck M, Kaping D, Marquez S, Everling S, Womelsdorf T. 2015. Mapping of Functionally Characterized Cell Classes onto Canonical Circuit Operations in Primate Prefrontal Cortex. Journal of Neuroscience. 35:2975–2991.

Avants BB, Tustison NJ, Song G, Cook PA, Klein A, Gee JC. 2011. A reproducible evaluation of ANTs similarity metric performance in brain image registration. NeuroImage. 54:2033– 2044.

Averbeck BB, Lee D. 2006. Effects of Noise Correlations on Information Encoding and Decoding. Journal of Neurophysiology. 95:3633–3644.

Belmonte JCI, Callaway EM, Caddick SJ, Churchland P, Feng G, Homanics GE, Lee K-F, Leopold DA, Miller CT, Mitchell JF, Mitalipov S, Moutri AR, Movshon JA, Okano H, Reynolds JH, Ringach DL, Sejnowski TJ, Silva AC, Strick PL, Wu J, Zhang F. 2015. Brains, Genes, and Primates. Neuron. 86:617–631.

Burman KJ, Palmer SM, Gamberini M, Rosa MGP. 2006. Cytoarchitectonic subdivisions of the dorsolateral frontal cortex of the marmoset monkey (Callithrix jacchus), and their projections to dorsal visual areas. J Comp Neurol. 495:149–172.

Carney HC, Hart E, Huk AC. 2019. Demonstration and quantification of memory-guided saccades in the common marmoset (with comparison to the macaque). Journal of Vision. 19:86a.

Constantinidis C, Goldman-Rakic PS. 2002. Correlated Discharges Among Putative Pyramidal Neurons and Interneurons in the Primate Prefrontal Cortex. Journal of Neurophysiology. 88:3487–3497.

Constantinidis C, Williams GV, Goldman-Rakic PS. 2002. A role for inhibition in shaping the temporal flow of information in prefrontal cortex. Nat Neurosci. 5:175–180.

Funahashi S. 2006. Prefrontal cortex and working memory processes. Neuroscience. 139:251– 261.

Funahashi S, Bruce CJ, Goldman-Rakic PS. 1989. Mnemonic coding of visual space in the monkey’s dorsolateral prefrontal cortex. Journal of Neurophysiology. 61:331–349.

Funahashi S, Bruce CJ, Goldman-Rakic PS. 1990. Visuospatial coding in primate prefrontal neurons revealed by oculomotor paradigms. Journal of Neurophysiology. 63:814–831.

Funahashi S, Bruce CJ, Goldman-Rakic PS. 1991. Neuronal activity related to saccadic eye movements in the monkey’s dorsolateral prefrontal cortex. Journal of Neurophysiology. 65:1464–1483.

Fuster JM. 1973. Unit activity in prefrontal cortex during delayed-response performance: neuronal correlates of transient memory. Journal of Neurophysiology. 36:61–78.

Fuster JM, Alexander GE. 1971. Neuron Activity Related to Short-Term Memory. Science. 173:652–654.

Goldman PS, Rosvold HE. 1970. Localization of function within the dorsolateral prefrontal cortex of the rhesus monkey. Experimental Neurology. 27:291–304.

Goldman PS, Rosvold HE, Vest B, Galkin TW. 1971. Analysis of the delayed-alternation deficit produced by dorsolateral prefrontal lesions in the rhesus monkey. Journal of Comparative and Physiological Psychology. 77:212–220.

Goldman-Rakic PS. 1995. Cellular basis of working memory. Neuron. 14:477–485.

Johnston K, Ma L, Schaeffer L, Everling S. 2019. Alpha Oscillations Modulate Preparatory Activity in Marmoset Area 8Ad. J Neurosci. 39:1855–1866.

Johnston KD, Barker K, Schaeffer L, Schaeffer D, Everling S. 2018. Methods for chair restraint and training of the common marmoset on oculomotor tasks. Journal of Neurophysiology. 119:1636–1646.

Jun JJ, Steinmetz NA, Siegle JH, Denman DJ, Bauza M, Barbarits B, Lee AK, Anastassiou CA, Andrei A, Aydın Ç, Barbic M, Blanche TJ, Bonin V, Couto J, Dutta B, Gratiy SL, Gutnisky DA, Häusser M, Karsh B, Ledochowitsch P, Lopez CM, Mitelut C, Musa S, Okun M, Pachitariu M, Putzeys J, Rich PD, Rossant C, Sun W, Svoboda K, Carandini M, Harris KD, Koch C, O’Keefe J, Harris TD. 2017. Fully integrated silicon probes for high-density recording of neural activity. Nature. 551:232–236.

Kishi N, Sato K, Sasaki E, Okano H. 2014. Common marmoset as a new model animal for neuroscience research and genome editing technology. Develop Growth Differ. 56:53– 62.

Kondo T, Saito R, Otaka M, Yoshino-Saito K, Yamanaka A, Yamamori T, Watakabe A, Mizukami H, Schnitzer MJ, Tanaka KF, Ushiba J, Okano H. 2018. Calcium Transient Dynamics of Neural Ensembles in the Primary Motor Cortex of Naturally Behaving Monkeys. Cell Reports. 24:2191-2195.e4.

Kubota K, Niki H. 1971. Prefrontal cortical unit activity and delayed alternation performance in monkeys. Journal of Neurophysiology. 34:337–347.

Li X, Morgan PS, Ashburner J, Smith J, Rorden C. 2016. The first step for neuroimaging data analysis: DICOM to NIfTI conversion. Journal of Neuroscience Methods. 264:47–56.

Link SW. 1982. Correcting response measures for guessing and partial information. Psychological Bulletin. 92:469–486.

Liu C, Ye FQ, Yen CC-C, Newman JD, Glen D, Leopold DA, Silva AC. 2018. A digital 3D atlas of the marmoset brain based on multi-modal MRI. NeuroImage. 169:106–116.

Lundqvist M, Rose J, Herman P, Brincat SL, Buschman TJ, Miller EK. 2016. Gamma and Beta Bursts Underlie Working Memory. Neuron. 90:152–164.

MacDonald SE, Pang JC, Gibeault S. 1994. Marmoset (Callithrix jacchus jacchus) spatial memory in a foraging task: Win-stay versus win-shift strategies. Journal of Comparative Psychology. 108:328–334.

Miles RC. 1957. Delayed-response learning in the marmoset and the macaque. Journal of Comparative and Physiological Psychology. 50:352–355.

Mitchell JF, Leopold DA. 2015. The marmoset monkey as a model for visual neuroscience. Neuroscience Research. 93:20–46.

Nakako T, Murai T, Ikejiri M, Ishiyama T, Taiji M, Ikeda K. 2013. Effects of a dopamine D1 agonist on ketamine-induced spatial working memory dysfunction in common marmosets. Behavioural Brain Research. 249:109–115.

Niki H, Watanabe M. 1976. Prefrontal unit activity and delayed response: Relation to cue location versus direction of response. Brain Research. 105:79–88.

Okano H, Hikishima K, Iriki A, Sasaki E. 2012. The common marmoset as a novel animal model system for biomedical and neuroscience research applications. Seminars in Fetal and Neonatal Medicine. 17:336–340.

Okano H, Sasaki E, Yamamori T, Iriki A, Shimogori T, Yamaguchi Y, Kasai K, Miyawaki A. 2016. Brain/MINDS: A Japanese National Brain Project for Marmoset Neuroscience. Neuron. 92:582–590.

Ott T, Nieder A. 2016. Dopamine D2 Receptors Enhance Population Dynamics in Primate Prefrontal Working Memory Circuits. Cereb Cortex. cercor;bhw244v1.

Paxinos G, Watson C, Petrides M, Rosa M, Tokuno H. 2012. The marmoset brain in stereotaxic coordinates.London; Waltham, MA: Academic Press.

Quintana J, Yajeya J, Fuster JM. 1988. Prefrontal representation of stimulus attributes during delay tasks. I. Unit activity in cross-temporal integration of sensory and sensory-motor information. Brain Research. 474:211–221.

Rao SG, Williams GV, Goldman-Rakic PS. 1999. Isodirectional Tuning of Adjacent Interneurons and Pyramidal Cells During Working Memory: Evidence for Microcolumnar Organization in PFC. Journal of Neurophysiology. 81:1903–1916.

Reser DH, Burman KJ, Yu H-H, Chaplin TA, Richardson KE, Worthy KH, Rosa MGP. 2013. Contrasting Patterns of Cortical Input to Architectural Subdivisions of the Area 8 Complex: A Retrograde Tracing Study in Marmoset Monkeys. Cerebral Cortex. 23:1901– 1922.

Riley MR, Constantinidis C. 2016. Role of Prefrontal Persistent Activity in Working Memory. Front Syst Neurosci. 9.

Sadoun A, Rosito M, Fonta C, Girard P. 2019. Key periods of cognitive decline in a nonhuman primate model of cognitive aging, the common marmoset (Callithrix jacchus). Neurobiology of Aging. 74:1–14.

Sasaki E. 2015. Prospects for genetically modified non-human primate models, including the common marmoset. Neuroscience Research. 93:110–115.

Sasaki E, Suemizu H, Shimada A, Hanazawa K, Oiwa R, Kamioka M, Tomioka I, Sotomaru Y, Hirakawa R, Eto T, Shiozawa S, Maeda T, Ito M, Ito R, Kito C, Yagihashi C, Kawai K, Miyoshi H, Tanioka Y, Tamaoki N, Habu S, Okano H, Nomura T. 2009. Generation of transgenic non-human primates with germline transmission. Nature. 459:523–527.

Selvanayagam J, Johnston KD, Schaeffer DJ, Hayrynen LK, Everling S. 2019. Functional Localization of the Frontal Eye Fields in the Common Marmoset Using Microstimulation. J Neurosci. 39:9197–9206.

Selvanayagam J, Wong RK, Everling S. 2021. Marmtouch: Experimental control for touchscreen experiments using a raspberry pi.

Spinelli S, Ballard T, Feldon J, Higgins GA, Pryce CR. 2006. Enhancing effects of nicotine and impairing effects of scopolamine on distinct aspects of performance in computerized attention and working memory tasks in marmoset monkeys. Neuropharmacology. 51:238–250.

Spinelli S, Pennanen L, Dettling AC, Feldon J, Higgins GA, Pryce CR. 2004. Performance of the marmoset monkey on computerized tasks of attention and working memory. Cognitive Brain Research. 19:123–137.

Sreenivasan KK, D’Esposito M. 2019. The what, where and how of delay activity. Nat Rev Neurosci. 20:466–481.

Trainito C, von Nicolai C, Miller EK, Siegel M. 2019. Extracellular Spike Waveform Dissociates Four Functionally Distinct Cell Classes in Primate Cortex. Current Biology. 29:2973-2982.e5.

Tsujimoto S, Sawaguchi T. 2002. Working memory of action: a comparative study of ability to selecting response based on previous action in New World monkeys (Saimiri sciureus and Callithrix jacchus). Behavioural Processes. 58:149–155.

Vijayraghavan S, Everling S. 2021. Neuromodulation of Persistent Activity and Working Memory Circuitry in Primate Prefrontal Cortex by Muscarinic Receptors. Front Neural Circuits. 15:648624.

Wang X-J. 2021. 50 years of mnemonic persistent activity: quo vadis? Trends in Neurosciences. 44:888–902.

Wilson FA, O’Scalaidhe SP, Goldman-Rakic PS. 1994. Functional synergism between putative gamma-aminobutyrate-containing neurons and pyramidal neurons in prefrontal cortex. Proc Natl Acad Sci USA. 91:4009–4013.

Yamazaki Y, Saiki M, Inada M, Watanabe S, Iriki A. 2016. Sustained performance by common marmosets in a delayed matching to position task with variable stimulus presentations. Behavioural Brain Research. 297:277–284.

