## Supplementary figures and images for "Delay-related activity in marmoset prefrontal cortex"

### Supplementary Figure 1

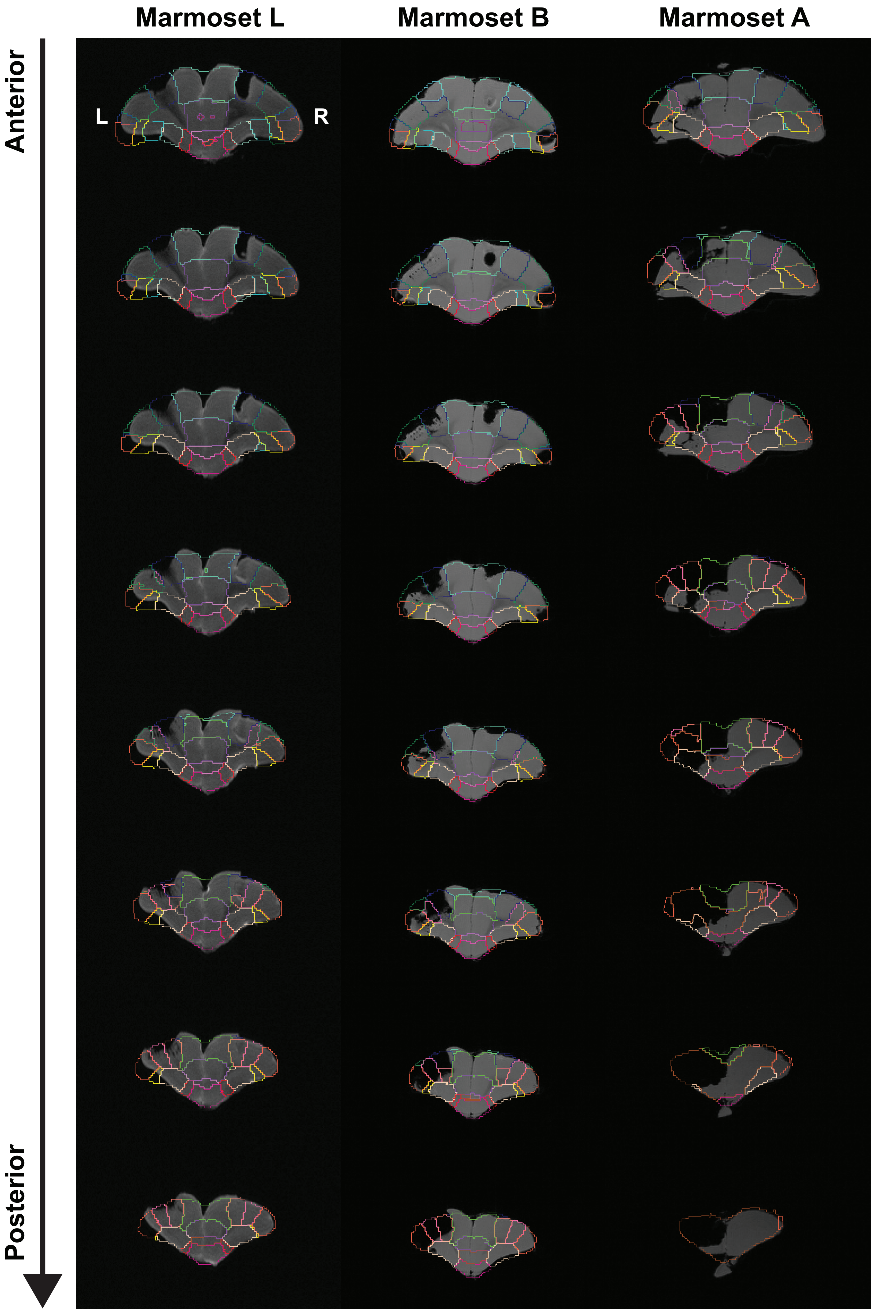

### Supplementary Figure 2

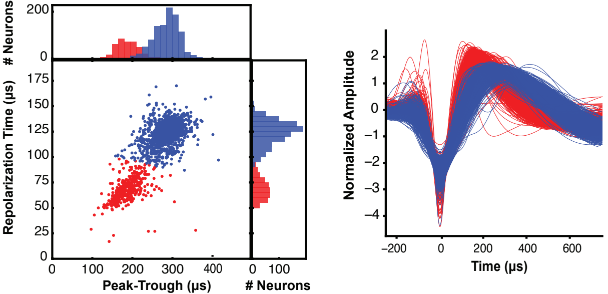
